# Structure of a tripartite protein complex that targets toxins to the type VII secretion system

**DOI:** 10.1101/2023.07.21.550046

**Authors:** Timothy A. Klein, Prakhar Y. Shah, Polyniki Gkragkopoulou, Dirk W. Grebenc, Youngchang Kim, John C. Whitney

## Abstract

Type VII secretion systems are membrane-embedded nanomachines used by Gram-positive bacteria to export effector proteins from the cytoplasm to the extracellular environment. Many of these effectors are polymorphic toxins comprised of an N-terminal Leu-x-Gly (LXG) domain of unknown function and a C-terminal toxin domain that inhibits the growth of bacterial competitors. In recent work, it was shown that LXG effectors require two cognate Lap proteins for T7SS-dependent export. Here, we present the 2.6Å structure of the LXG domain of the TelA toxin from the opportunistic pathogen *Streptococcus intermedius* in complex with both of its cognate Lap targeting factors. The structure reveals an elongated α-helical bundle within which each Lap protein makes extensive hydrophobic contacts with either end of the LXG domain. Remarkably, despite low overall sequence identity, we identify striking structural similarity between our LXG complex and PE-PPE heterodimers exported by the distantly related ESX type VII secretion systems of Mycobacteria implying a conserved mechanism of effector export among diverse Gram-positive bacteria. Overall, our findings demonstrate that LXG domains, in conjunction with their cognate Lap targeting factors, represent a tripartite secretion signal for a widespread family of T7SS toxins.

## Introduction

The type VII secretion system (T7SS) is an integral membrane protein complex that facilitates the export of protein effectors through the cell envelope of Gram-positive bacteria. Likely owing to their distinct cell envelope architectures, bacteria belonging to the phyla Actinobacteria and Firmicutes encode distinct T7SS complexes referred to as T7SSa and T7SSb, respectively, with each subtype possessing several phylum-specific subunits (1). The ESX-1 T7SSa is well-studied in pathogenic Mycobacteria such as *Mycobacterium tuberculosis* because it is a major virulence factor required for the phagosomal escape step that precedes cytoplasmic replication of infected macrophages (2). Conversely, the T7SSb is less well understood but has recently been shown to play a role in both virulence and/or bacterial antagonism in both pathogenic and environmental bacteria (3–7).

The T7SSb is known to export two families of effector proteins. WXG100 proteins, which include the ubiquitous T7SS effector EsxA, are approximately 100 amino acid long α-helical proteins whose precise function is unclear but are unique in that they are required for overall T7SS apparatus function and are also secreted from the cell (8, 9). Besides these small non-enzymatic effectors, large multi-domain polymorphic toxins are also exported by several characterized T7SSb pathways with the best studied examples belonging to the LXG toxin family (4). LXG toxins contain a loosely conserved N-terminal leucine-x-glycine (LXG) domain from which the protein family derives its name and diverse C-terminal toxin domains (10). LXG toxins from several T7SSb-expressing bacteria including *Bacillus subtilis*, *Staphylococcus aureus*, *Enterococcus faecalis*, and *Streptococcus intermedius* have been identified and in most instances, the toxin domains have been shown to possess antibacterial activities that mediate bacterial antagonism (4–6, 11). While recent work has advanced our mechanistic understanding of how some of these toxins inhibit bacterial growth, less is known about the role of their N-terminal LXG domains (3, 4, 11). These domains have long been hypothesized to target effectors to the membrane embedded T7SSb apparatus, however, this has yet to been shown experimentally (10).

Although the function of the LXG domain is incompletely understood, our group recently demonstrated that some of the toxins harboring these domains require so-called targeting factors for their export (12). These targeting factors, termed <u>L</u>XG-associated <u>α</u>-helical <u>p</u>roteins <u>1</u> and <u>2</u> (Lap1 and Lap2), directly interact with LXG domains and are necessary for LXG toxin export. As is often the case for bacterial gene products that physically interact, *lap* genes co-occur with the gene encoding their cognate LXG toxin and are typically found upstream of the toxin gene. Lap targeting factors are small α-helical proteins that are structurally reminiscent of WXG100 effectors. Interestingly, Lap1 proteins possess a conserved FxxxD motif similar to the YxxxD/E export motif of T7SSa effectors (13, 14). Furthermore, mutation of this motif in the LapD1 targeting factor abrogates secretion of its cognate LXG toxin, TelD (12).

In the present study, we identify two additional families of targeting factors, termed Lap3 and Lap4, and show that members of these families found upstream of the *S. intermedius* LXG-containing toxin TelA are required for effector secretion. Using X-ray crystallography, we solve the structure of these targeting factors in complex with the LXG domain of TelA, revealing the molecular basis of how LXG domains simultaneously interact with both of their targeting factors. Using TelA as a model, we subsequently show that the LXG domain is both necessary and sufficient for protein export by the T7SSb and identify key amino acid motifs involved in this process. Finally, using Alphafold2 models informed by our structure, we provide evidence that diverse LXG toxin complexes adopt a conserved tripartite architecture at their N-terminus that is required for their secretion by the T7SSb.

## Results

### Identification of two new families of LXG toxin targeting factors

While many LXG toxins are encoded downstream of a Lap1-Lap2 pair (formerly DUF3130-DUF3958), others are found downstream of DUF5344 and DUF5082 encoding genes, which are also predicted to encode for small α-helical proteins (Fig S1) (8). The opportunistic pathogen *Streptococcus intermedius* B196 (Si^B196^) has a single *lap1*-*lap2* associated effector, TelC, and two DUF5344-DUF5082 associated effectors, TelA and TelB (Fig 1A). Because of their genetic association to LXG effectors and their predicted α-helical secondary structure, we henceforth refer to DUF5344-DUF5082 pairs as Lap3 and Lap4 to reflect their predicted similarity to Lap1 and Lap2, respectively (Fig S1). This similarity also led us to hypothesize that like *lap1* and *lap2* deletion strains, strains lacking either *lap3* or *lap4* would be unable to export their downstream effector. To test this prediction, we focused on the TelA toxin and its corresponding Lap proteins, LapA3 and LapA4, to examine the role of these uncharacterized protein families in LXG toxin export. In line with our hypothesis, Si^B196^ strains bearing inactivating mutations in either *lapA3* or *lapA4* fail to export TelA and this secretion defect could be restored through plasmid-borne expression of the deleted *lap* genes in the corresponding deletion strains (Fig 1B).

**Figure 1.**
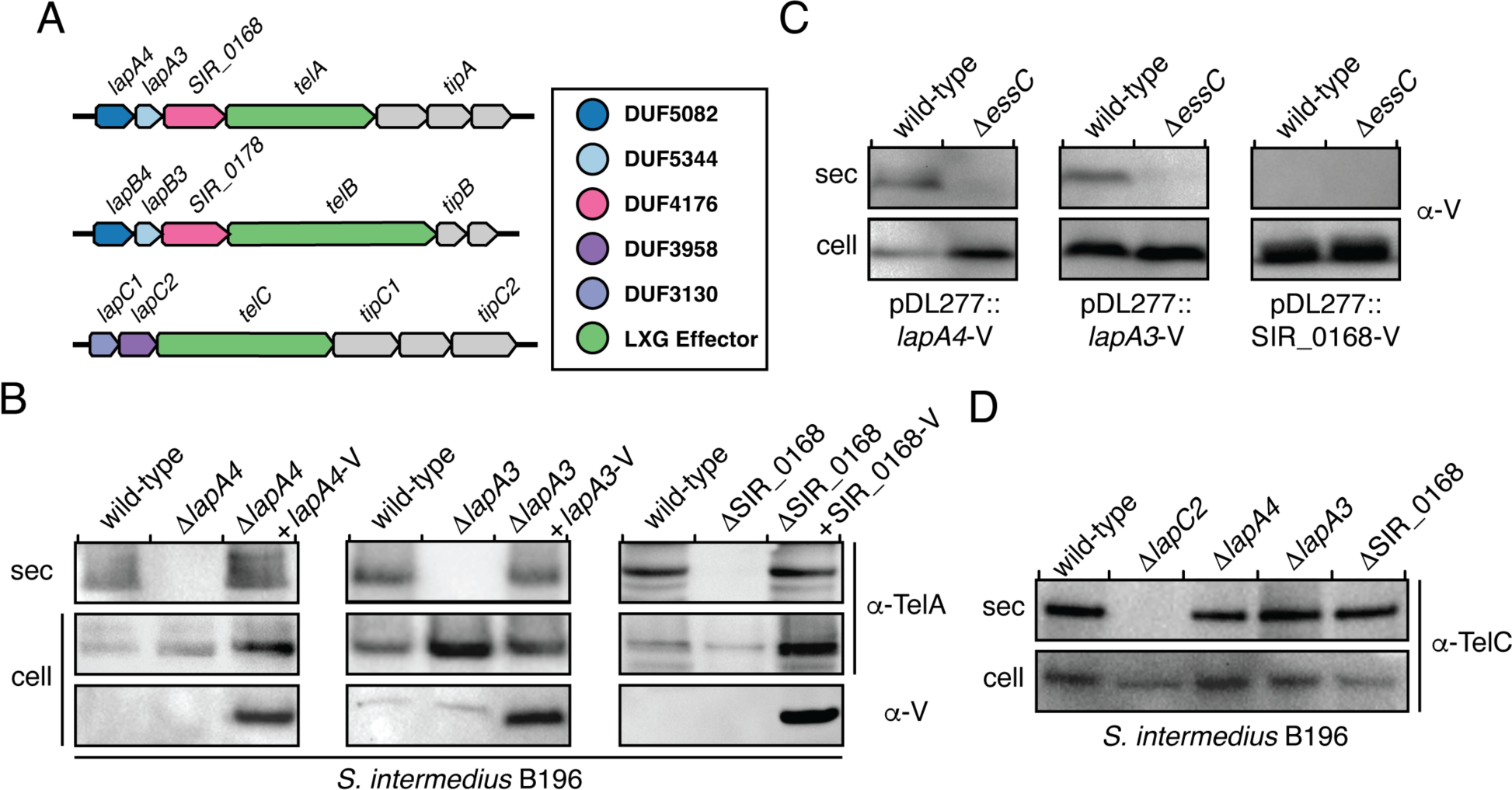
| TelA toxin is co-secreted with its cognate targeting factors. A) Genomic context of the genes encoding the three Tel toxins exported by the T7SS of *S. intermedius* B196, their cognate Tip immunity proteins, and Lap targeting factors. B) Western blot showing that *lapA4*, *lapA3*, and *SIR_0168* are required for the secretion of TelA toxin. C) LapA4 and LapA3 are exported from cells in a T7SS-dependent manner whereas SIR_0168 is not. D) Lap targeting factors are specific for their cognate Tel toxin.

The *telA* and *telB* gene neighborhoods possess an additional gene that encodes for a member of the DUF4176 family of uncharacterized proteins, which are absent in the *telC* gene cluster (Fig 1A). To test the potential involvement of DUF4176 proteins in LXG toxin secretion, we generated a strain lacking the DUF4176 encoding gene upstream of *telA*, SIR_0168, and again assayed for TelA export. As is the case for *lapA3* and *lapA4* deficient strains of Si^B196^, TelA is not secreted by a strain lacking SIR_0168 and secretion could be restored by trans complementation of SIR_0168 (Fig 1B).

We next wanted to test if the protein product of each gene required for TelA secretion is exported from cells. We and others previously found that the targeting factors for TelC toxins, LapC1 and LapC2, are not secreted from cells (12, 15). To test if this is also the case for LapA3 and LapA4, we expressed C-terminal VSV-G tagged versions of these proteins and performed western blot analyses on cell and culture supernatant fractions of wild-type and T7SS-null (ι1*essC*) strains of Si^B196^. In contrast to LapC1 and LapC2, we could readily detect T7SSb-dependent export of LapA3 and LapA4 (Fig 1C). We also found that the Lap3-Lap4 pair associated with the TelB LXG toxin, LapB3 and LapB4, are similarly secreted in a T7SSb-dependent manner (Fig S2). This difference in localization for the different Lap pairs suggests that Lap3 and Lap4 proteins may promote the secretion of their cognate effectors in a way that is mechanistically distinct from Lap1-Lap2. Finally, we also tested for secretion of VSV-G tagged SIR_0168 and found that this protein is not secreted by Si^B196^ indicating that despite being required for TelA secretion, it is likely not exported from cells alongside TelA, LapA3, and LapA4 (Fig 1C).

WXG100 effectors such as EsxA are typically encoded next to genes encoding the T7SSb apparatus and based on structural studies of T7SSa systems, are likely involved in the export of all T7SS effectors due to their role in the formation of an active T7SS apparatus pore (16–18). By contrast, each *tel* gene in the Si^B196^ genome is found adjacent to a unique set of *lap* genes. This observation led us to hypothesize that Lap proteins are highly specific and are therefore likely only required for the secretion of their cognate effector. To test this, we used a TelC-specific antibody to assay for cellular and secreted levels of TelC in our *lapA3*, *lapA4*, and SIR_0168-deficient strains. The deletion of any of the three genes required for TelA secretion had no effect on extracellular levels of TelC whereas a strain lacking the LapC2 targeting factor was unable to export TelC (Fig 1D). Taken together, these data indicate that the targeting factors encoded upstream of LXG genes are indeed effector specific and are therefore not required for global T7SS effector export.

### Structure of the TelA_LXG_-LapA3-LapA4 complex

To better understand the function of LXG domains and their potential interactions with Lap targeting factors, we next sought to determine their three-dimensional structures by X-ray crystallography. We previously showed that TelC, LapC1, and LapC2 physically interact to form a heteromeric complex, however, crystallization of this particle proved refractory (12). Therefore, we adopted a similar co-expression and co-purification strategy with the LXG domain of TelA (residues 1-224) or TelB (residues 1-219) along with their cognate targeting factors and found that like TelC they interact to form a 1:1:1 complex (Fig 2A and Fig S3A and S3B). Of these two complexes, TelA_LXG_-LapA3-LapA4 crystallized readily, and we were able to solve its crystal structure to a resolution of 2.6Å (Table 1).

**Figure 2.**
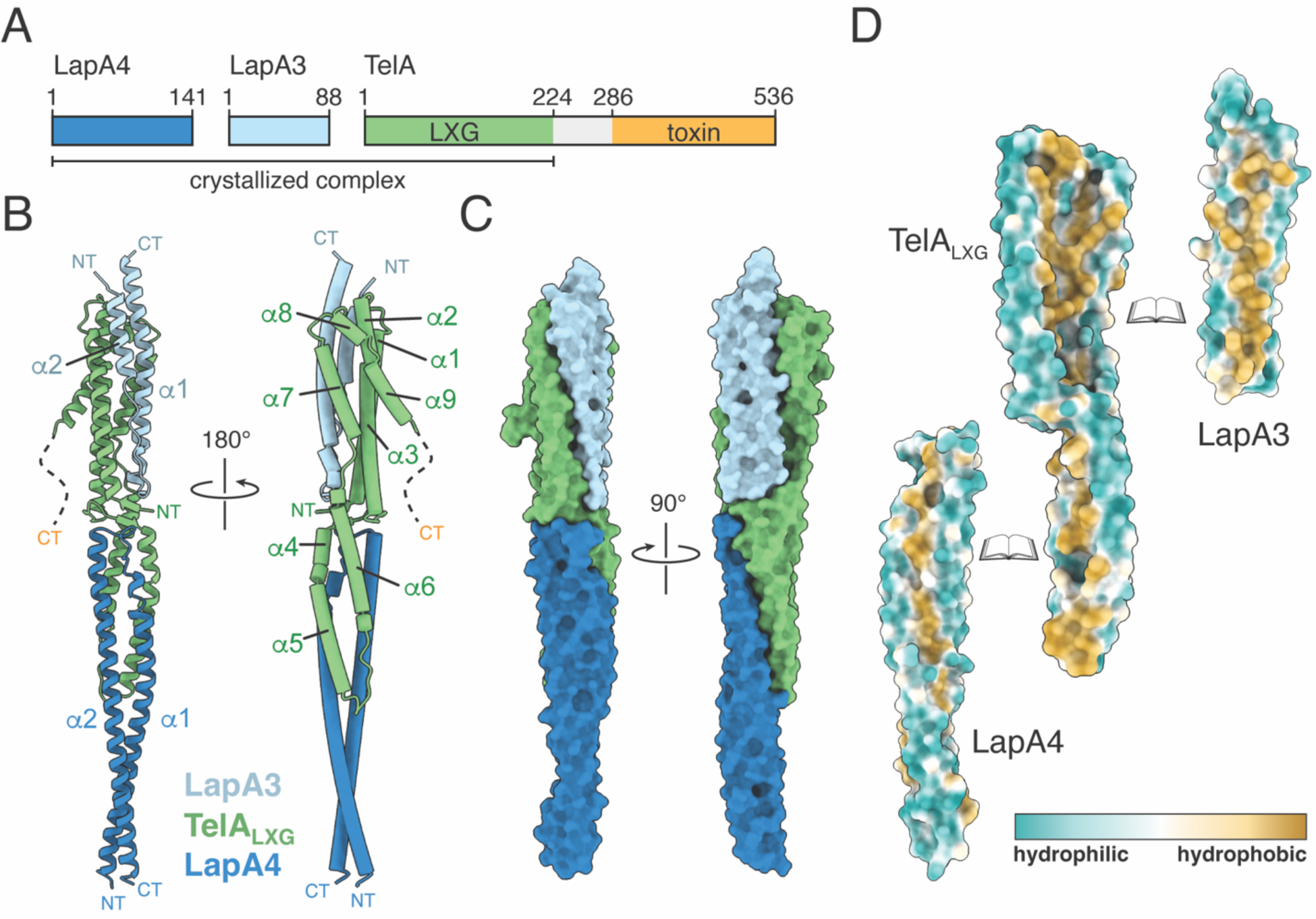
| Overall structure of the TelA_LXG_-LapA4-LapA3 complex. A) Domain schematic of LapA4, LapA3, and TelA depicting the amino acid boundaries of the crystallized complex. B) Cartoon model of the TelA_LXG_-LapA4-LapA3 complex shown from two opposing views. The secondary structure elements and termini of each protein subunit are indicated. The approximate location of TelA’s middle region (residues 224-286) and C-terminal toxin domain (CT) is indicated by a dashed line. C) Space-filling model of the TelA_LXG_-LapA4-LapA3 complex shown from two orthogonal angles. D) Hydrophobic surface representation depicting the interaction interfaces between TelA_LXG_ and each of its Lap targeting factors.

**Table 1.** X-ray data collection and refinement statistics.11.

|  | TelA_1-223:LapA3:LapA4 (native) |
| --- | --- |
| **Data Collection** |  |
| Wavelength (Å) | 0.9791897918 |
| Space group | P1 |
| Cell dimensions |  |
| *a*, *b*, *c* (Å) | 51.9, 61.4, 97.9 |
| $\alpha$, $\beta$, $\gamma$ (°) | 77.7, 77.1, 65.1 |
| Resolution<sup>a</sup> (Å) | 45.08-2.60 (2.644-2.60) |
| Unique reflections | 31528 ((1504) |
| CC<sub>1/2</sub><sup>c</sup> | 0.948 (0.478) |
| *R*<sub>merge</sub><sup>b</sup> | 0.209 (0.858) |
| *R*<sub>pim</sub><sup>c</sup> | 0.123 (0.586) |
| *I*/σ*I* | 77.9 (11.0) |
| Completeness (%) | 97.2 (92.9) |
| Redundancy | 3.7 (2.7) |
| Wilson B-factor | 41.5 |
| **Refinement** |  |
| *R*<sub>work</sub><sup>d</sup> / *R*<sub>free</sub> (%) | 21.55/25.8 |
| Average B-factors (Å<sup>2</sup>) |  |
| Protein | 5656.2 |
| Water/Other | 4747.5/6060.9 |
| No. atoms |  |
| Protein | 6614 |
| Water/Other | 275/12 |
| Rms deviations |  |
| Bond lengths (Å) | 00.001 |
| Bond angles (°) | 00.300 |
| Ramachandran plot (%) |  |
| Total favored<sup>e</sup> | 9898.84 |
| Total allowed | 11.16 |
| PDB code | 8GMH |

<sup>a</sup>Values in parentheses correspond to the highest resolution shell. <sup>b</sup> $R_{\text{merge}} = \sum_h \sum_j |I_{hj} - \langle I_h \rangle| / \sum_h \sum_j I_{hj}$ , where *I*<sub>hj</sub> is the intensity of observation *j* of reflection *h*. <sup>c</sup>As defined by Karplus and Diederichs (52). <sup>d</sup> $R = \sum_h |F_o| - |F_c| / \sum_h |F_o|$ for all reflections, where *F*<sub>o</sub> and *F*<sub>c</sub> are observed and calculated structure factors, respectively. *R*<sub>free</sub> is calculated analogously for the test reflections, randomly selected, and excluded from the refinement. <sup>e</sup>As defined by Molprobit(53)

The overall structure of TelA_LXG_-LapA3-LapA4 reveals that the complex formed by these proteins forms an elongated α-helical bundle with approximate dimensions of ∼180Å by ∼30Å (Fig 2B). Within this bundle structure, LapA3 and LapA4 both adopt a helix-turn-helix topology with the α-helices of the former aligning approximately parallel to one another whereas those of the latter cross over at the protein’s termini. TelA_LXG_ is comprised of nine α-helices that are divided into two sub-domains by a central region containing two short antiparallel β-strands (Fig 2C). Each of these sub-domains interacts with a single Lap protein with one sub-domain consisting of α-helices 4-6 and interacting with LapA4 and the other comprising α-helices 1-3 and 7-9 and interacting with LapA3. Remarkably, despite the relatively small size of each of the subunits within this complex, the buried surface areas between TelA_LXG_-LapA3 and TelA_LXG_-LapA4 are 2261.5Å^2^ and 1810.4Å^2^, respectively. Analysis of the surface properties of these interfaces reveals that they are highly hydrophobic in nature suggesting that complex formation is largely entropically driven (Fig 2D).

### The TelA_LXG_ and LapA3 subcomplex resembles substrates of Mtb ESX T7SSs

Upon initial inspection of the TelA_LXG_-LapA3-LapA4 structure it was immediately apparent that it possesses many similarities to structurally characterized T7SSa effectors including EspB and the PE25-PPE41 heterodimer exported by the ESX-1 and ESX-5 systems of *M. tuberculosis*, respectively (Fig 3A)(14, 19). EspB and PE25-PPE41 heterodimers adopt similar all α-helical folds with the former resembling a fused version of the latter. These two T7Ssa effectors share a similar shape and topology with the TelA_LXG_ and LapA3 components of our T7SSb effector complex. Unlike TelA, EspB and PE25-PPE41 do not harbour C-terminal toxin domains and instead are proposed to oligomerize into pores in the outer membrane of *M. tuberculosis* (20, 21). Given that T7SSb-containing Firmicutes lack an outer membrane, it is perhaps not surprising that we observe no such oligomeric arrangement for TelA_LXG_-LapA3-LapA4 in the crystal lattice of our structure.

**Figure 3.**
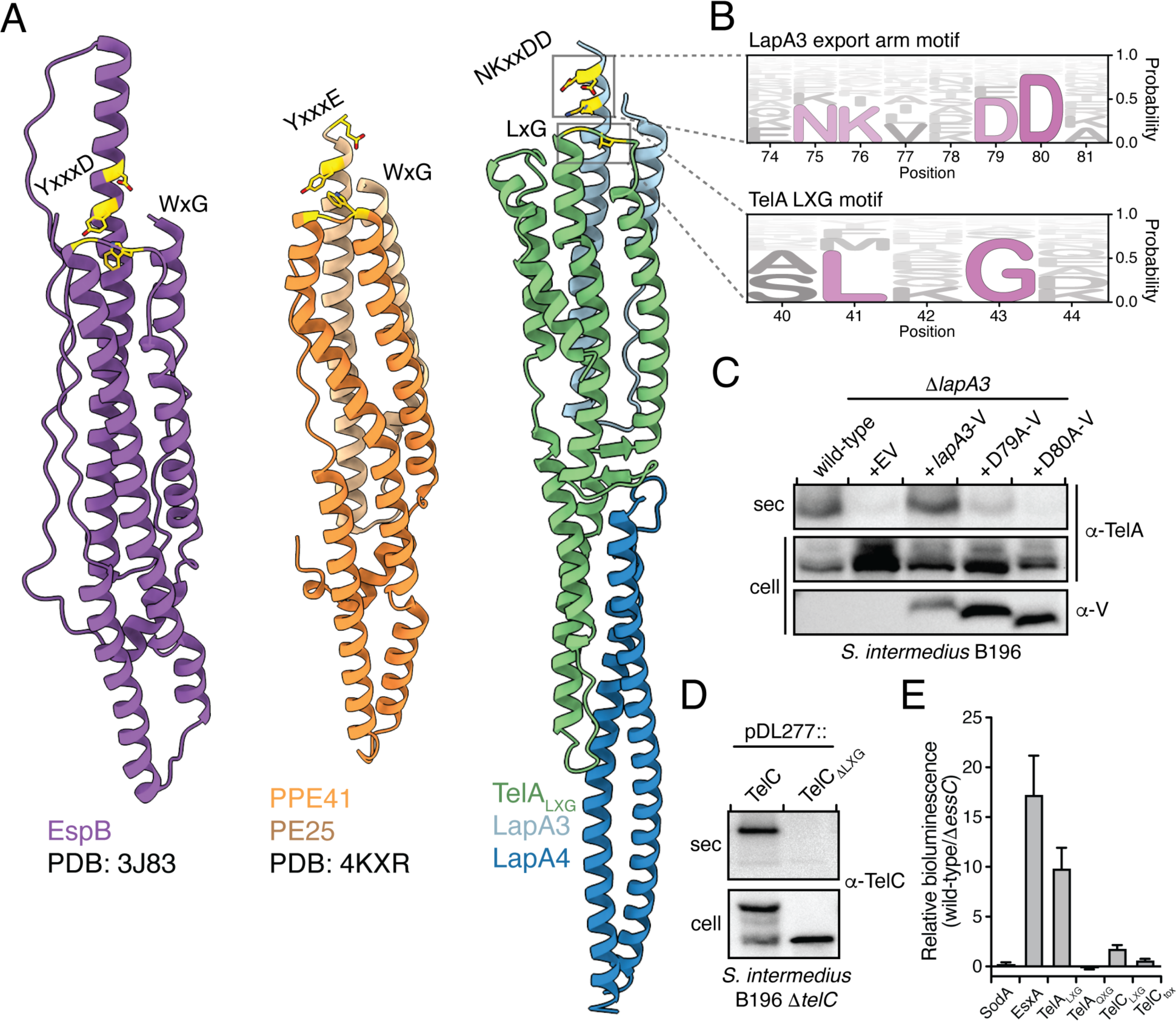
| LXG complexes are necessary and sufficient for T7SSb toxin export. A) Structural comparison between TelA_LXG_-LapA4-LapA3 complex and the well-characterized T7SSa substrates EspB (PDB code 3J83) and PE25-PPE41 (PDB code 4KXR) from *M. tuberculosis*. Regions highlighted yellow indicated the residues comprising the conserved export arm in the *M. tuberculosis* substrates and the comparable residues in TelA and LapA3. B) Sequence logo representation illustrating the conservation of the export arm and LXG motifs of LapA3 and TelA, respectively. C) Conserved LapA3 export arm motif residues are required for TelA toxin secretion from *S. intermedius* B196. D) The LXG domain of TelC is required for its export via the T7SSb. Western blot analysis of the cell and secreted (sec) fractions of *S. intermedius* B196 strains expressing full-length TelC or TelC lacking its N-terminal LXG domain (TelC_ΔLXG_). E) LXG domains of Tel effectors along with their cognate Lap proteins are sufficient for type VII secretion. Split luciferase assay in which the indicated proteins were fused to the small subunit of nanoluciferase and the bioluminescence of culture supernatants containing purified nanoluciferase large subunit was measured. Superoxide dismutase (SodA) was used as a cytoplasmic protein control. For additional details, see Experimental Procedures.

We next used multiple sequence alignments to search for conserved motifs within TelA_LXG_, LapA3, and LapA4 to identify residues that are likely important for function. Not surprisingly, given our prior analyses of TelC_LXG_, Lap1, and Lap2 sequences, we found that these proteins contain little overall sequence conservation (12). Nonetheless, two prominent motifs emerged from our analysis with the first being TelA’s LxG motif from which LXG toxins derive their name and the second being an NKxxDD motif found at the C-terminus of LapA3 (Fig 3A). Interestingly, these motifs are found near one another in three-dimensional space and bear a striking resemblance to the well-characterized export arm motif of T7SSa substrates (13, 14, 22). For example, Leu41 of the LxG motif is found in the same turn region of TelA as is the conserved WxG motif of PPE41. In both proteins, the conserved hydrophobic residue in these motifs is buried in a hydrophobic pocket of its interacting partner to buttress its solvent exposed C-terminal α-helix. The second motif is less well conserved between T7Ssa and T7SSb substrates in that the tyrosine residue found in the YxxxD/E export arm motif of EspB and PE25 is absent in LapA3. Nevertheless, all three proteins possess the conserved solvent exposed acidic residue within this motif in approximately the same position in their structure with LapA3 uniquely possessing two aspartic acid residues at this site (Fig 3B). Given the importance of residues in this position for type VII secretion, we mutated each conserved aspartic acid residue of LapA3 to alanine and examined the ability of these variants to facilitate TelA secretion. In line with their high degree of sequence conservation, expression of the D79A variant resulted in a substantial reduction in TelA secretion whereas the levels of secreted TelA in cells expressing the D80A variant were reduced to that of the Δ*lapA3* negative control (Fig 3C). When taken together with mutagenesis analysis of other T7SS substrates, these data indicate that a conserved C-terminal acidic residue appears to be a universal requirement for protein export by the T7SS apparatus (9, 12, 13).

### LXG complexes target toxins for export by the T7SSb

Our data so far suggest that LXG-Lap-Lap heterotrimeric complexes represent a tripartite trafficking complex that targets LXG toxins for T7SSb-dependent export from the cell. To test this hypothesis, we first examined the ability of TelC to be exported from cells in the absence of its LXG domain (TelC_ι1LXG_). We chose TelC for this initial experiment because our antibody for this protein recognizes its toxin domain whereas our TelA antibody recognizes TelA’s LXG domain. In line with our prediction, we found that in contrast to full-length TelC, which is readily exported from cells, TelC_ι1LXG_ remains in the cytoplasm (Fig 3D). We next sought to examine if LXG complexes are sufficient for T7SSb-dependent protein export. To test this, we employed a recently developed split luciferase assay in which an 11 amino acid fragment of deep-sea shrimp luciferase is fused to a protein of interest and that protein’s export is assessed by measuring the ability of the fused fragment to combine with the remainder of the luciferase enzyme added exogenously via its bioluminescence activity (23). After confirming the suitability of this assay for use in Si^B196^ using EsxA export as a positive control, we fused the luciferase fragment to the LXG domains of TelA and TelC, co-expressed these domains with their cognate Lap proteins, and measured whether these heterotrimeric complexes are sufficient for protein export via the T7SSb. Consistent with functioning as a T7SSb targeting complex, we could readily detect TelA and TelC LXG domain export despite these proteins lacking their C-terminal toxin domains (Fig 3E). Furthermore, we tested the functional importance of the conserved leucine residue that comprises the LXG motif by making a single amino acid substitution to glutamine in TelA_LXG_ (TelA_QXG_). In line with playing a critical role in the export process, we were unable to detect luminescence from the culture supernatant of a strain expressing TelA_QXG_ heterotrimeric complexes. Importantly, SDS-PAGE analysis of purified TelA_QXG_ -LapA3-LapA4 complex demonstrates that this mutation does not affect complex formation and therefore instead likely plays a role in targeting TelA to the T7SSb apparatus (Fig S3C). Collectively, these data indicate that LXG-domain containing complexes are both necessary and sufficient for type VIIb secretion.

### LXG targeting complexes adopt a conserved architecture

While the structure and function of TelA’s toxin domain is unknown and its Alphafold2 predicted structure is low confidence, we reasoned that we could use our experimental structure of TelA_LXG_-LapA3-LapA4 to predict the structure of other full-length LXG toxins for which the structure of the toxin has been determined or is confidently predicted (Fig 4). Like TelA, we found AF2 predictions of TelB to be low confidence likely owing to the presence of a long linker region connecting its N-terminal LXG domain to its C-terminal toxin domain (4). By contrast, we were able to generate high confidence models of full-length TelC because an X-ray structure is available for its toxin domain, and of full-length YxiD toxin from *B. subtilis* because it lacks a middle linker region and possesses a confidently predicted nuclease toxin (4, 5). Based on these results and preliminary structural data described by others, we propose that full-length toxin-Lap-Lap complexes adopt a tomahawk shaped structure with the metaphorical blade and shaft subunits being formed by the toxin domain and LXG-Lap-Lap complex, respectively(24). Given that there are currently over 4000 annotated LXG toxins in publicly available sequence databases, we expect that our structure will allow for the high confidence modelling of the vast majority of tripartite LXG secretion complexes found in diverse species of T7SS-containing Firmicutes.

**Figure 4.**
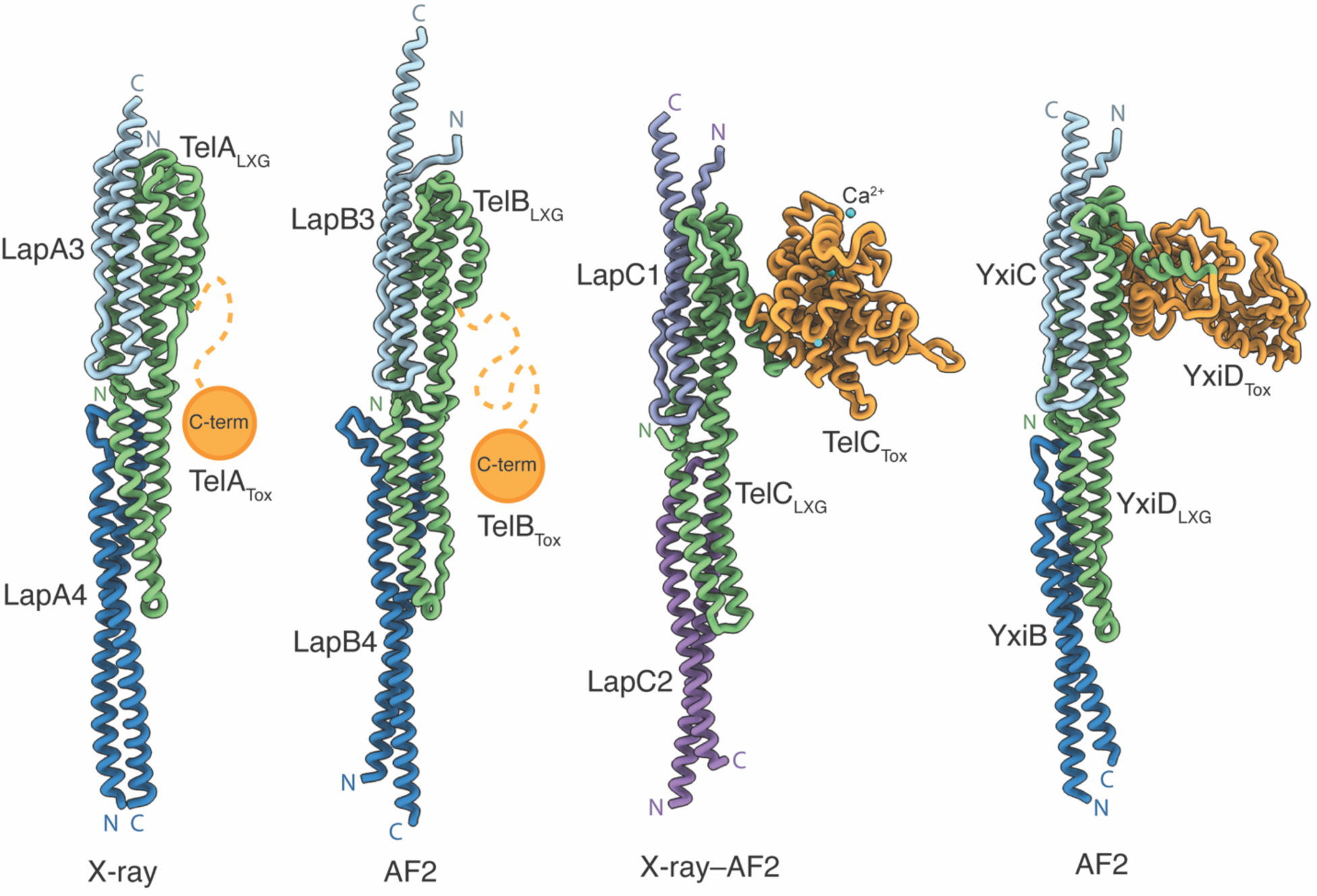
| LXG toxin complexes adopt a conserved tripartite architecture at their N-terminus and harbor diverse C-terminal toxin domains. Experimental and Alphafold2 predicted structures of LXG toxins and their cognate Lap targeting factors. Low confidence toxin domains are depicted as orange circles (TelA and TelB) whereas experimentally determined (TelC_tox_, PDB code 5UKH) and high confidence toxin domains (YxiD_tox_ from *B. subtilis*) are shown as orange ribbons.

## Discussion

In this study, we have determined the first structure of an LXG domain and demonstrated that it represents the minimum requirement for LXG effector export by the T7SSb apparatus. In addition, we find that targeting factors belonging to the DUF5082 and DUF5344 protein families are co-exported as part of an N-terminal toxin trafficking complex required for type VIIb secretion. Given the widespread distribution of these protein families among Firmicutes, we anticipate our findings will be broadly applicable to LXG toxins from diverse bacterial species.

While our work defines the minimal requirement for type VIIb secretion, in some instances, additional cellular factors are required for LXG toxin export. For example, we found that TelA export also requires the DUF4176 family member, SIR_0168. While SIR_0168 itself is not secreted from cells, we speculate that it may function as a cytosolic chaperone that maintains TelA-LapA3-LapA4 complexes in a secretion competent state, perhaps in a manner that is analogous to EspG proteins from T7SSa systems (19, 25). However, it is unclear why some effectors require a DUF4176 protein whereas others do not. One possibility is that LXG toxins with an extended middle domain between their LXG and toxin domains, like TelA and TelB, require a DUF4176 protein for intracellular trafficking whereas those that lack this middle domain, like TelC, do not (4). The site of action of the TelA and TelB toxins is in the cytoplasm of competitor bacteria whereas TelC acts extracellularly. Therefore, one tantalizing possibility is that the middle domain is involved in toxin translocation across the membrane of target cells and that the function of DUF4176 proteins in toxin-producing cells is to prevent erroneous membrane insertion prior to toxin export via the T7SSb.

The asymmetric structural models of LXG-Lap-Lap complexes raises interesting questions regarding the folded state of LXG toxins as they translocate the T7SSb apparatus. While recent cryo-electron microscopy structures of the ESX-1 and ESX-5 T7SSa apparatuses show the closed conformation of the EccC/EssC translocase, estimates of the dimensions of the open complex based on structural comparisons to the related ATPase FtsK suggest its diameter is approximately 30Å, which closely matches that of the N-terminal shaft complex comprised of the LXG domain and its two cognate targeting factors (17, 18, 26, 27). This observation leads us to speculate that the C-terminal toxin domains likely exist in an unfolded state as they transit the T7SSb apparatus. Otherwise, their asymmetric shape would not be conducive to export through the symmetrical pore of the T7SS apparatus because doing so would likely compromise essential ion gradients required for cellular viability. Furthermore, toxin domain unfolding prior to export would also provide a plausible mechanism for immunity protein dissociation in toxin-producing bacteria.

Another outstanding question that remains is how LXG toxin complexes are specifically recognized by the T7SSb apparatus. A likely candidate to fulfill this role is a C-terminal region of EssC that consists of a portion of its second ATPase domain and the entirety of its third ATPase domain (28, 29). In contrast to the rest of the T7SSb apparatus components, several bioinformatic studies show that the amino acid sequence of this region of EssC varies significantly even between strains of the same bacterium and likely coevolved with a suite of cognate LXG toxins (28, 30). While our attempts to observe a physical interaction between purified Tel pre-secretion complexes and various regions of EssC have thus far been met with limited success, this is perhaps not surprising as it remains unclear how the conformational states of the EssC translocon are regulated and such regulatory inputs may be needed before EssC and LXG toxin complexes can physically interact. Based on studies of T7SSa systems, these inputs likely include interaction with other components of the T7SSb apparatus, EsxA induced multimerization, and ATP hydrolysis by the ATPase domains of EssC (16, 18). Future investigations into the specific roles that each of these components plays in the export process will be critical to furthering our understanding of the molecular mechanism of type VII effector secretion.

## EXPERIMENTAL PROCEDURES

### Bacterial strains, plasmids, and growth conditions

All gene knockouts were introduced into a *S. intermedius* B196 wild-type background and genomic DNA from this organism was used as a PCR template for all molecular cloning. *E. coli* XL1-Blue and BL21 (DE3) CodonPlus were used for molecular cloning and protein overexpression, respectively. The complete list of bacterial strains used in this study are listed in Table S1. pET29b- and pETDuet-1-derived plasmids were used for recombinant protein expression in *E. coli*, while pDL277-derived plasmids were used for constitutive protein expression in *S. intermedius*. PCR amplification of *S. intermedius* genes was done with Phusion polymerase (NEB). PCR amplicons were digested with NcoI/SalI for pETDuet-1 MCS1, NdeI/XhoI for pET29b/pETDuet-1 MCS2, or BamHI/SalI for pDL277 and the genes of interest were ligated into the respective plasmids with T4 DNA ligase (NEB). Additionally, some primers included an additional 3′ sequence to generate tagged proteins with poly-6-His (HHHHHH), VSV-G (YTDIEMNRLGK) or GSG-linker-pep86 (GSGVSGWRLFKKIS) for protein purification, western blot detection, or nano-luciferase based secretion assays, respectively. When cloning into pDL277, the P96 promoter sequence described by Lo Sapio and colleagues from *Streptococcus pneumoniae* was fused upstream to the gene of interest using <u>s</u>plicing by <u>o</u>verlap <u>e</u>xtension (SOE) PCR (31). A comprehensive list of the plasmids used in this study are found in Table S2. *E. coli* was grown in lysogeny broth at 37°C at 225 revolutions per minute under aerobic conditions. The growth media was supplemented with 50 μL/mL of kanamycin, 150 μL/mL of carbenicillin, or 100 μL/mL spectinomycin to select for pET29b, pETDuet-1, or pDL277, respectively. *S. intermedius* was grown in Todd Hewitt broth supplemented with 0.5% yeast extract (THY) in a 37°C stationary 5% CO_2_ incubator. All *S. intermedius* strains were grown on THY agar plates for 1-3 days prior to growth in THY broth to ensure uniform growth rate. THY broth and agar plates were supplemented with 250 μL/mL of kanamycin or 75 μL/mL of spectinomycin to select for gene deletions and pDL277-derived plasmids, respectively.

### DNA manipulation

To prepare genomic DNA, *S. intermedius* overnight cultures were pelleted and resuspended in a 10:1 ratio of culture to InstaGene Matrix (BioRad) after which the extraction protocol as per manufacturer instructions was followed. The DNA primers used in this study were generated by Integrated DNA Technology (IDT). Molecular cloning was done using Phusion polymerase, common restriction endonucleases, and T4 DNA ligase from NEB. Sanger sequencing was performed by Azenta Life Sciences and The Center for Applied Genomics at The Hospital for Sick Children in Toronto, Ontario.

### Transformation of *S. intermedius*

Linear DNA fragments for allelic replacement (see below) or pDL277-derived plasmids were transformed into *S. intermedius* as previously described (32). In brief, *S. intermedius* overnight cultures were back diluted 1:10 into 2 mL THY broth supplemented with 5 μL of 0.1 mg/mL *S. intermedius* competence stimulating peptide (DSRIRMGFDFSKLFGK, synthesized by Genscript) and incubated for 2 hours at the appropriate growth conditions (see above). 100-200ng of linear or plasmid DNA was then added, and cultures were grown for an additional 3 hours before plating on THY agar plates supplemented with the appropriate antibiotic.

### Gene deletion in *S. intermedius* by allelic exchange

*S. intermedius* gene deletions were performed as described previously (33). In short, gene deletion constructs were assembled in pETDuet-1 plasmid using SOE PCR. These constructs were comprised of 1000bps upstream of the gene of interest (5′ flank) including the first 15-45bps of the gene of interest ORF, a spectinomycin promoter derived from pDL277, a kanamycin resistance cassette from pBAV1K, and 1000bps downstream of the gene of interest (3′ flank) including the last 15-45bps of the gene of interest ORF (34). These final deletion constructs had the following generic arrangement: pETduet-1::5′-flank_SpecPromoter_kanR_3′-flank and were subsequently digested from the plasmid using BamHI and NotI (NEB). The resultant insert (5′-flank_SpecPromoter_kanR_3′-flank) was gel extracted (Monarch DNA GEL Extraction Kit, NEB) and added to competence peptide stimulated *S. intermedius* cells (see above).

### Secretion assays

Culture tubes containing 2 mL THY broth were inoculated with *S. intermedius* colonies from THY agar plates and grown overnight (final OD_600_ of 1.0-1.1). Overnight cultures were then back diluted 1:100 into fresh 20 mL THY and grown overnight for a second iteration. Cell and secreted (sec) fractions were then separated by centrifugation at 4,000 *g* for 15 minutes. The cell samples were washed once in 1 mL of PBS pH 7.4 and centrifuged at 4,000 *g* for 5 minutes. PBS wash was decanted, and the cell samples were resuspended in 150 μL of 1:1 ratio of PBS to 4x Laemmli buffer, after which the cell samples were boiled for 10 minutes. The decanted sec samples were treated with trichloroacetic acid to a 10% final volume and the samples were incubated at 4°C overnight to precipitate secreted protein. The sec samples were then centrifuged at 35,000 *g* for 30 minutes and resulting pellets were washed once with 20 mL of 95% cold acetone. Sec samples were centrifuged again at 35,000 *g* for 30 minutes and the acetone was removed. The sec samples were allowed to air dry briefly on ice in a fume hood. The precipitated protein pellets were then resuspended in 100 μL of 1:2 ratio of 4x Laemmli buffer to 8M urea and boiled for 10 minutes. Cell and sec samples were analyzed for proteins of interest by SDS-PAGE and western blotting.

### Nano-luciferase assay

Nano-luciferase (NanoLuc) based secretion assay was performed as described by Yang and colleagues (23). In brief, 2 mL THY cultures were inoculated with *S. intermedius* colonies from a THY agar plate and grown overnight to an OD_600_ of 1.0-1.1. Cultures were centrifuged at 4000 *g* for 10 minutes and 100 μL of supernatant was aliquoted into clear bottom, black walled 96-well plate (Corning #3631). NanoLuc large subunit 11S was purified and added into the reaction well at a final concentration of 5 μM as per published protocol (35). 2 μL of furimazine substrate was added to the reaction well from a working reagent stock of a 2:100 NanoDLR Stop & Glo Substrate to Buffer ratio (Promega #N1610). A total well volume of approximately 112.5 μL was incubated at room temperature for 2 minutes prior to measuring luminescence and OD_600_ readings on an EnVision plate reader (PerkinElmer).

### Antibody generation

A custom polyclonal antibody for TelA was generated for this study. Because full-length TelA toxin is toxic to *E. coli*, overexpression of just the LXG domain of TelA (TelA_LXG_) was performed for antibody production (4). To improve TelA_LXG_ stability, we co-expressed and co-purified TelA_LXG_ with LapA4. This complex was purified in PBS pH 7.4 and yielded 10 mg of total protein that was subsequently shipped to Genscript for custom antibody production. The generation of the α-TelC antibody was described previously (4).

### SDS-PAGE, SYPRO red staining, and western blotting

A tris-tricine buffering system (200mM Tris, 100mM Tricine, 0.1% SDS, pH 8.3) was used for all SDS-PAGE gels run in this study to better differentiate proteins less than 30 kDa (36). In-gel imaging of proteins was done using SYPRO Red protein gel stain (Invitrogen). The gels were rinsed in deionized water, stained in 1:5000 SYPRO red, 10% acetic acid for one hour, destained in 7.5% acetic acid for 1 minute and imaged using a ChemiDoc imaging system (Bio-Rad). For western blotting, gels were wet transferred to nitrocellulose at 100V for 30 minutes (α-TelC and α-V short for α-VSV-G) or to methanol-activated PVDF at 80V for 1 hour (α-EsxA). The blots were blocked with 5% skim milk in TBS-T for 30 minutes before being incubated with primary antibody for 1 hour (1:5000 titer for α-TelC and α-EsxA, and 1:3000 titer for α-V). Blots were washed three times with 15 mL of TBS-T and then incubated with α-rabbit secondary antibody (1:5000 titer) for 45 minutes. Blots were washed three more times in TBS-T before being developed with Clarity Max Western ECL reagent and imaged on a ChemiDoc XRS+ (Bio-Rad).

### Protein expression and purification

Wild-type and mutant TelA_LXG_-LapA3-LapA4 protein complexes were expressed in BL21 (DE3) CodonPlus. The strains were back diluted 1:50 in LB supplemented with 50 μL/mL of kanamycin and 150 μL/mL of carbenicillin and grown to an OD_600_ 0.50-0.55. Protein expression was then induced by the addition of 1 mM IPTG and protein was expressed overnight (∼18 hours) at 18°C. The cells were pelleted, resuspended in lysis buffer (20 mM Tris-HCl pH 8.0, 300 mM NaCl, 10 mM imidazole), and sonicated four times for 30 seconds at 30% amplitude. Cellular debris was cleared by 30 minutes of centrifugation at 35,000 *g*. The lysate sample was run on a benchtop Ni-NTA column and washed thrice with 20 mL of lysis buffer. Protein was eluted with 4 mL of elution buffer (20 mM Tris-HCl pH 8.0, 300 mM NaCl, 400 mM imidazole). For protein crystallization trials, the eluants were then further purified by size exclusion chromatography using a HiLoad 16/600 Superdex 200 or Superdex 200 Increase 10/300 GL columns hooked up to a ÄKTA explorer (Cytiva). A similar protocol was used for the purification of TelB_LXG_-LapB3-LapB4 complex except that lysis and elution buffers were additionally supplemented with 1mM dithiothreitol.

### Protein crystallization

Native TelA_LXG_-LapA3-LapA4 complex at 10 mg/ml was screened for crystallization conditions using the hanging drop vapor diffusion method and the MCSG suite of sparse matrix crystal screens (Anatrace). Crystals formed after one to two weeks in 0.02M MgCl_2_, 0.1M HEPES:NaOH pH 7.5, and 22% (w/v) polyacrylic acid 5100. The crystals were cryoprotected using the same buffer supplemented with 25% ethylene glycol and flash frozen in liquid nitrogen prior to X-ray data collection.

### X-ray data collection, structure determination, and model refinement

Data collection was performed using the Structural Biology Center sector 19-ID beamline at the Advanced Photon Source. Diffraction data were collected at 100K with 0.3s exposure and a 0.5 degree of rotation for a total of 400 degrees. The crystal used to solve the structure diffracted to 2.6 Å and diffraction images were collected on a Dectris Pilatus 3 X 6M detector using an X-ray wavelength of 12.662 keV (0.97918 Å). Diffraction data were processed using HKL3000 software and the structure of TelA_LXG_-LapA3-LapA4 was solved by molecular replacement using MOLREP implemented in HKL3000 with an AlphaFold2 model as the search model (37, 38). Coot was used to adjust the model manually to the electron density while computational structural refinement was performed with Phenix.refine until the *R*_work_ and *R*_free_ converged to 21.5% and 25.8%, respectively (39, 40). The final model includes two copies of each protein in the unit cell of space group P1. All collection and refinement statistics for TelA_LXG_-LapA3-LapA4 can be found in Table 1 and the structure is deposited in the Protein Data Bank under PDB code 8GMH.

### Protein structure prediction, visualization, and analysis

Protein secondary structure predictions were generated by PSIPRED 4.0 on the UCL PSIPRED Workbench at hyperlink: http://bioinf.cs.ucl.ac.uk/psipred/ (41). Protein complex predictions were performed with Colabfold AlphaFold2 using MMseqs2 locally installed on an HPE Apollo 6500 system running Red Hat Enterprise Linux with Nvidia Quadro RTX 8000 GPUs (42, 43). Predicted models were manually assessed and selected based on the per-residue pLDDT scores for each chain and the biological plausibility of the predicted complex. UCSF ChimeraX was used for protein structure analysis and figure generation (44). Surface hydrophobicity calculations were performed using default parameters (45). Interchain interfaces were analyzed using PDBsum (46). All structural alignments and reported RMSD scores were calculated by the DALI webserver using PDB search and Pairwise alignment tools (47).

### Sequence analysis and sequence logo generation

Homologous sequences to LapA3, LapA4, and TelA_LXG_ were identified using JackHMMER (HmmerWeb v2.41.2) searches of the UniprotKB database, restricted to the phylum Firmicutes, iterating until at least 150 sequences were obtained (48). Accessions were downloaded and full sequences of active entries were subsequently retrieved from Uniprot. Duplicate sequences were removed and the remaining aligned using ClustalO with default parameters (49). HMMs for each sequence alignment were generated using the Skylign webserver, set to “create HMM – remove mostly empty columns” (50). The resulting matrices were downloaded as tabular text, formatted, and then visualized using Logomaker (51).

### SEC-MALS analysis of protein complexes

Size exclusion chromatography (SEC) with multi-angle laser light scattering (MALS) was performed on TelA_LXG_-LapA3-LapA4 and TelB_LXG_-LapB3-LapB4 protein complexes. The proteins were expressed and purified as described above and concentrated to 2 mg/ml by spin filtration. SEC was run on a Superdex 200 column (GE Healthcare), and MALS was conducted using a MiniDAWN and Optilab system (Wyatt Technologies). Data was collected and analyzed using the Astra software package (Wyatt Technologies).

## DATA AVAILABILITY

The X-ray structure factors for the TelA_LXG_-LapA3-LapA4 complex have been deposited in the Protein Data Bank under the accession number 8GMH. All data generated or analysed during this study are available from the corresponding authors upon request.

## AUTHOR CONTRIBUTIONS

T.A.K. and J.C.W. conceived the study. All authors contributed to experimental design. T.A.K., P.Y.S., and P.G. generated strains and plasmids. T.A.K. and P.Y.S. performed protein expression, purification, and crystallization. T.A.K., D.W.G., and Y.K. solved and analyzed the crystal structure. T.A.K., P.Y.S., and P.G. performed biochemical experiments. T.A.K., P.Y.S., and J.C.W. analyzed the data. T.A.K., P.Y.S., and J.C.W. wrote the paper. All authors provided feedback on the manuscript.

## ACKNOWLEDGEMENTS

The authors thank Tracy Palmer for sharing protocols and reagents, Giuseppe Melacini and Madoka Akimoto for access to and training on the SEC-MALS system, and members of the Whitney Lab for helpful discussions. We sincerely thank the members of the Structural Biology Center (SBC) at Argonne National Laboratory for their help with data collection at the 19-ID beamline. The use of SBC beamlines at the Advanced Photon Source is supported by the U.S. Department of Energy (DOE) Office of Science and operated for the DOE Office of Science by Argonne National Laboratory under contract no. DE-AC02-06CH11357. T.A.K. and P.Y.S. are supported by Canada Graduate Scholarships from the Natural Sciences and Engineering Research Council of Canada (NSERC). This work was supported by a project grant (PJT-173486) from the Canadian Institutes of Health Research (CIHR).

## REFERENCES

1. H. R. Tran, D. W. Grebenc, T. A. Klein, J. C. Whitney, Bacterial type VII secretion: An important player in host-microbe and microbe-microbe interactions. Molecular microbiology 115, 478–489 (2021).

2. A. M. Abdallah et al., Type VII secretion--mycobacteria show the way. Nature reviews. Microbiology 5, 883–891 (2007).

3. Z. Cao, M. G. Casabona, H. Kneuper, J. D. Chalmers, T. Palmer, The type VII secretion system of Staphylococcus aureus secretes a nuclease toxin that targets competitor bacteria. Nat Microbiol 2, 16183 (2016).

4. J. C. Whitney et al., A broadly distributed toxin family mediates contact-dependent antagonism between gram-positive bacteria. eLife 6 (2017).

5. K. Kobayashi, Diverse LXG toxin and antitoxin systems specifically mediate intraspecies competition in Bacillus subtilis biofilms. PLoS genetics 17, e1009682 (2021).

6. A. Chatterjee, J. L. E. Willett, G. M. Dunny, B. A. Duerkop, Phage infection and sub-lethal antibiotic exposure mediate Enterococcus faecalis type VII secretion system dependent inhibition of bystander bacteria. PLoS genetics 17, e1009204 (2021).

7. M. L. Burts, W. A. Williams, K. DeBord, D. M. Missiakas, EsxA and EsxB are secreted by an ESAT-6-like system that is required for the pathogenesis of Staphylococcus aureus infections. Proceedings of the National Academy of Sciences of the United States of America 102, 1169–1174 (2005).

8. L. Bowman, T. Palmer, The Type VII Secretion System of Staphylococcus. Annual review of microbiology 75, 471–494 (2021).

9. T. A. Sysoeva, M. A. Zepeda-Rivera, L. A. Huppert, B. M. Burton, Dimer recognition and secretion by the ESX secretion system in Bacillus subtilis. Proceedings of the National Academy of Sciences of the United States of America 111, 7653–7658 (2014).

10. D. Zhang, R. F. de Souza, V. Anantharaman, L. M. Iyer, L. Aravind, Polymorphic toxin systems: Comprehensive characterization of trafficking modes, processing, mechanisms of action, immunity and ecology using comparative genomics. Biology direct 7, 18 (2012).

11. F. R. Ulhuq et al., A membrane-depolarizing toxin substrate of the Staphylococcus aureus type VII secretion system mediates intraspecies competition. Proceedings of the National Academy of Sciences of the United States of America 10.1073/pnas.2006110117 (2020).

12. T. A. Klein et al., Dual Targeting Factors Are Required for LXG Toxin Export by the Bacterial Type VIIb Secretion System. mBio 10.1128/mbio.02137-22, e0213722 (2022).

13. M. H. Daleke et al., General secretion signal for the mycobacterial type VII secretion pathway. Proceedings of the National Academy of Sciences of the United States of America 109, 11342–11347 (2012).

14. M. Solomonson et al., Structure of EspB from the ESX-1 type VII secretion system and insights into its export mechanism. Structure 23, 571–583 (2015).

15. W. K. Teh et al., Characterization of TelE, a T7SS LXG Effector Exhibiting a Conserved C-Terminal Glycine Zipper Motif Required for Toxicity. Microbiol Spectr 10.1128/spectrum.01481-23, e0148123 (2023).

16. O. S. Rosenberg et al., Substrates Control Multimerization and Activation of the Multi-Domain ATPase Motor of Type VII Secretion. Cell 161, 501–512 (2015).

17. N. Famelis et al., Architecture of the mycobacterial type VII secretion system. Nature 576, 321–325 (2019).

18. C. M. Bunduc et al., Structure and dynamics of a mycobacterial type VII secretion system. Nature 593, 445–448 (2021).

19. N. Korotkova et al., Structure of the Mycobacterium tuberculosis type VII secretion system chaperone EspG5 in complex with PE25-PPE41 dimer. Molecular microbiology 94, 367–382 (2014).

20. J. Piton, F. Pojer, S. Wakatsuki, C. Gati, S. T. Cole, High resolution CryoEM structure of the ring-shaped virulence factor EspB from Mycobacterium tuberculosis. J Struct Biol X 4, 100029 (2020).

21. Q. Wang et al., PE/PPE proteins mediate nutrient transport across the outer membrane of Mycobacterium tuberculosis. Science 367, 1147–1151 (2020).

22. Y. Lou, J. Rybniker, C. Sala, S. T. Cole, EspC forms a filamentous structure in the cell envelope of Mycobacterium tuberculosis and impacts ESX-1 secretion. Molecular microbiology 103, 26–38 (2017).

23. Y. Yang, F. Alcock, H. Kneuper, T. Palmer, A high throughput assay to measure Type VII secretion in Staphylococcus aureus. bioRxiv 10.1101/2023.06.03.543475, 2023.2006.2003.543475 (2023).

24. Y. Yang et al., Three small partner proteins facilitate the type VII-dependent secretion export of an antibacterial nuclease. bioRxiv 10.1101/2023.04.01.535202, 2023.2004.2001.535202 (2023).

25. M. H. Daleke et al., Specific chaperones for the type VII protein secretion pathway. The Journal of biological chemistry 287, 31939–31947 (2012).

26. M. Zoltner et al., EssC: domain structures inform on the elusive translocation channel in the Type VII secretion system. The Biochemical journal 473, 1941–1952 (2016).

27. N. Poweleit et al., The structure of the endogenous ESX-3 secretion system. eLife 8 (2019).

28. B. Warne et al., The Ess/Type VII secretion system of Staphylococcus aureus shows unexpected genetic diversity. BMC genomics 17, 222 (2016).

29. F. Jager, H. Kneuper, T. Palmer, EssC is a specificity determinant for Staphylococcus aureus type VII secretion. Microbiology (Reading) 164, 816–820 (2018).

30. K. Bowran, T. Palmer, Extreme genetic diversity in the type VII secretion system of Listeria monocytogenes suggests a role in bacterial antagonism. Microbiology (Reading) 167 (2021).

31. M. Lo Sapio, M. Hilleringmann, M. A. Barocchi, M. Moschioni, A novel strategy to over-express and purify homologous proteins from Streptococcus pneumoniae. Journal of biotechnology 157, 279–286 (2012).

32. T. Tomoyasu et al., Role of catabolite control protein A in the regulation of intermedilysin production by Streptococcus intermedius. Infection and immunity 78, 4012–4021 (2010).

33. T. A. Klein et al., Structure of the Extracellular Region of the Bacterial Type VIIb Secretion System Subunit EsaA. Structure 10.1016/j.str.2020.11.002 (2020).

34. A. V. Bryksin, I. Matsumura, Rational design of a plasmid origin that replicates efficiently in both gram-positive and gram-negative bacteria. PloS one 5, e13244 (2010).

35. G. C. Pereira et al., A High-Resolution Luminescent Assay for Rapid and Continuous Monitoring of Protein Translocation across Biological Membranes. Journal of molecular biology 431, 1689–1699 (2019).

36. H. Schagger, Tricine-SDS-PAGE. Nature protocols 1, 16–22 (2006).

37. W. Minor, M. Cymborowski, Z. Otwinowski, M. Chruszcz, HKL-3000: the integration of data reduction and structure solution--from diffraction images to an initial model in minutes. Acta crystallographica. Section D, Biological crystallography 62, 859–866 (2006).

38. A. Vagin, A. Teplyakov, Molecular replacement with MOLREP. Acta crystallographica. Section D, Biological crystallography 66, 22–25 (2010).

39. P. Emsley, B. Lohkamp, W. G. Scott, K. Cowtan, Features and development of Coot. Acta crystallographica. Section D, Biological crystallography 66, 486–501 (2010).

40. P. V. Afonine et al., Towards automated crystallographic structure refinement with phenix.refine. *Acta crystallographica. Section D*, Biological crystallography 68, 352–367 (2012).

41. D. W. A. Buchan, D. T. Jones, The PSIPRED Protein Analysis Workbench: 20 years on. Nucleic acids research 47, W402–W407 (2019).

42. J. Jumper et al., Highly accurate protein structure prediction with AlphaFold. Nature 596, 583–589 (2021).

43. M. Mirdita et al., ColabFold: making protein folding accessible to all. Nature methods 19, 679–682 (2022).

44. T. D. Goddard et al., UCSF ChimeraX: Meeting modern challenges in visualization and analysis. Protein science : a publication of the Protein Society 27, 14–25 (2018).

45. B. Testa, P. A. Carrupt, P. Gaillard, F. Billois, P. Weber, Lipophilicity in molecular modeling. Pharmaceutical research 13, 335–343 (1996).

46. R. A. Laskowski, PDBsum: summaries and analyses of PDB structures. Nucleic acids research 29, 221–222 (2001).

47. L. Holm, Using Dali for Protein Structure Comparison. Methods Mol Biol 2112, 29–42 (2020).

48. R. D. Finn et al., HMMER web server: 2015 update. Nucleic acids research 43, W30–38 (2015).

49. F. Sievers et al., Fast, scalable generation of high-quality protein multiple sequence alignments using Clustal Omega. Mol Syst Biol 7, 539 (2011).

50. T. J. Wheeler, J. Clements, R. D. Finn, Skylign: a tool for creating informative, interactive logos representing sequence alignments and profile hidden Markov models. BMC bioinformatics 15, 7 (2014).

51. A. Tareen, J. B. Kinney, Logomaker: beautiful sequence logos in Python. Bioinformatics 36, 2272–2274 (2020).

52. P. A. Karplus, K. Diederichs, Linking crystallographic model and data quality. Science 336, 1030–1033 (2012).

53. I. W. Davis, L. W. Murray, J. S. Richardson, D. C. Richardson, MOLPROBITY: structure validation and all-atom contact analysis for nucleic acids and their complexes. Nucleic acids research 32, W615–619 (2004).

