## Supplemental Material for "Structure of a tripartite protein complex that targets toxins to the type VII secretion system"

**Running title**: Structure of the T7SSb export signal

Keywords: type VII secretion system, protein export, bacterial toxins

* To whom correspondence should be addressed: J.C.W

Telephone: (+1) 905-525-9140

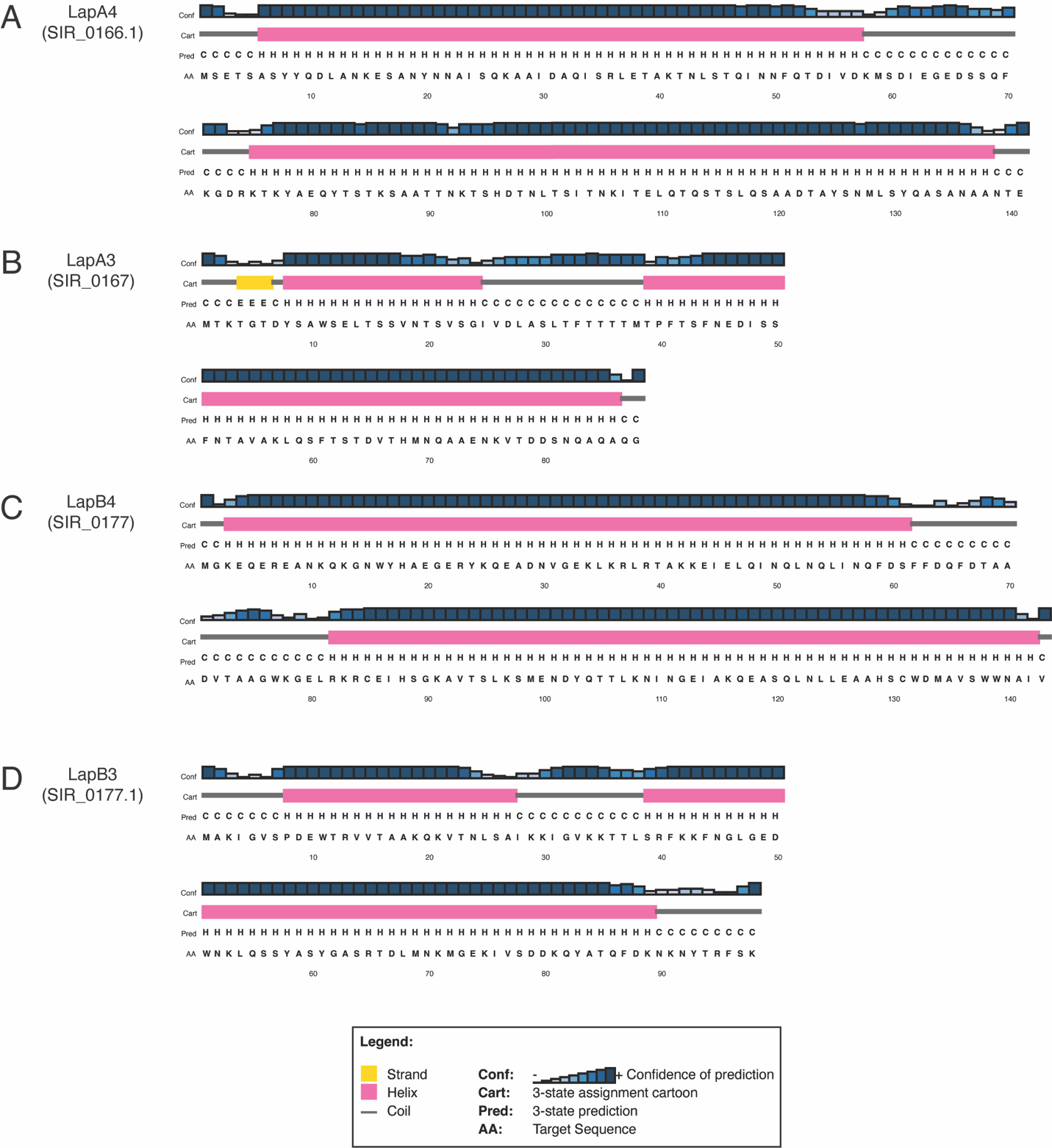

**Fig. S1 | Secondary structure predictions for the Lap3 and Lap4 proteins from *S. intermedius* B196.** Graphical output from PSIPRED 4.0 secondary structure analyses of **a,** LapA4. **b,** LapA3. **c,** LapB4. **d,** LapB3. Per-residue secondary structure predictions and confidence scores are indicated above each amino acid in the sequence as indicated in the legend.

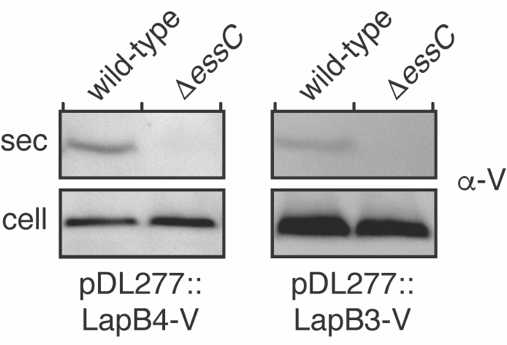

**Fig. S2 | LapB3 and LapB4 are exported from *S. intermedius* B196 in a T7SS-dependent manner.** Western blot analysis of the cell and supernatant (sec) fractions of *S. intermedius* B196 strains expressing VSV-G epitope tagged LapB4 (LapB4-V, left) or VSV-G epitope tagged LapB3 (LapB3-V, right).

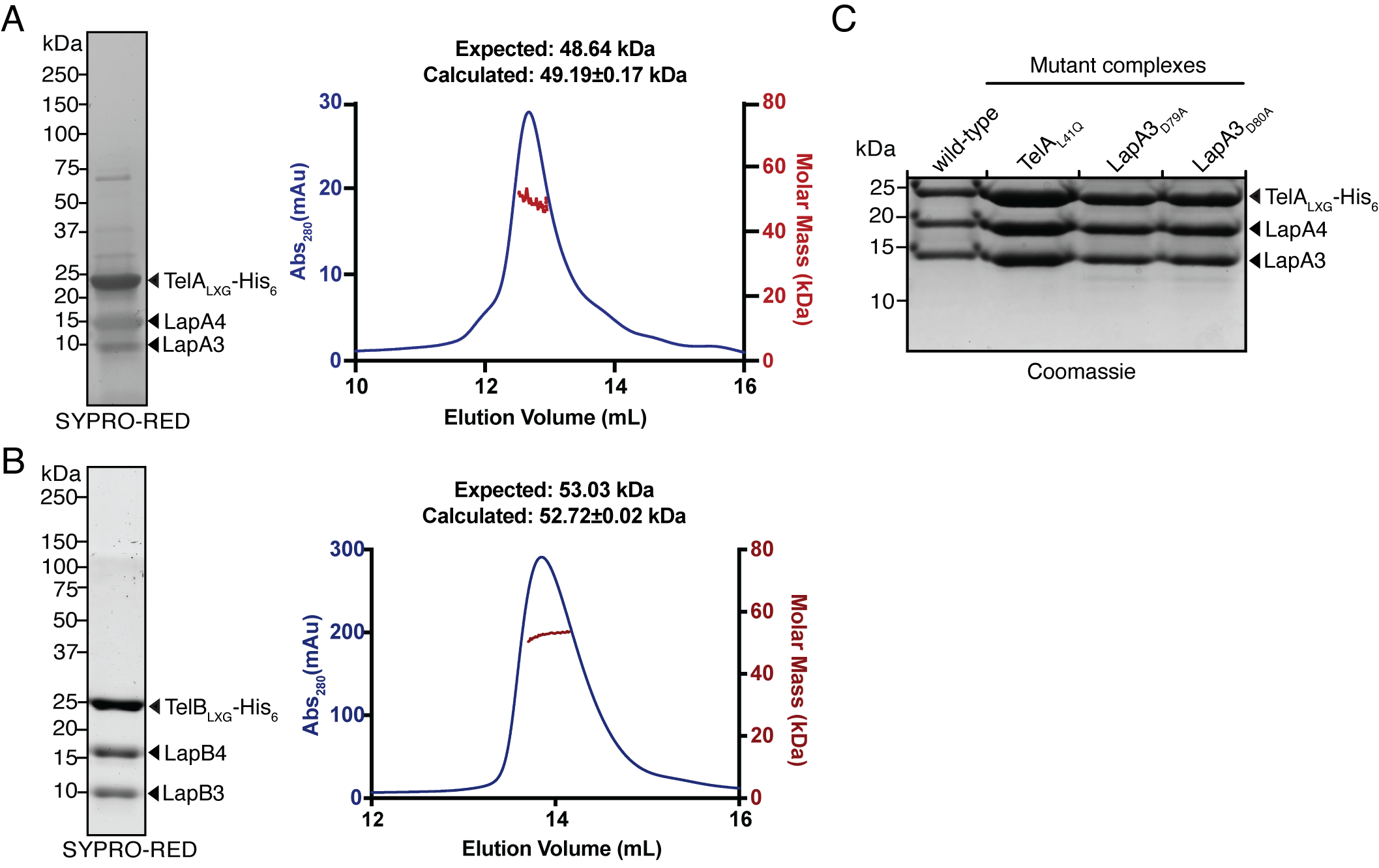

**Fig. S3 | TelA_LXG_-LapA4-LapA3 and TelB_LXG_-LapB4-LapB3 form heterotrimeric complexes with 1:1:1 stoichiometry.** **a,b,** SDS-PAGE and SEC-MALS analysis of purified TelA_LXG_-LapA4-LapA3 and TelB_LXG_-LapB4- LapB3 complexes. Expected molecular weight values represent the sum of the molecular weights of each protein in each complex. **c,** Mutagenesis of conserved residues in TelA and LapA3 does not impact heterotrimer formation *in vitro*. SDS-PAGE analysis of co-purified wild-type TelA_LXG_-LapA4-LapA3 and the indicated site-specific variants.

**Table S1: Strains used in this study.**

| **Organism** | **Genotype** | **Description** | **Reference** |
| --- | --- | --- | --- |
| *S. intermedius* B196 | Wildtype |  | (1) |
|  | ΔSIR_0166.1::kan^R^ | *lapA4* deletion strain | This study |
|  | ΔSIR_0167::kan^R^ | *lapA3* deletion strain | This study |
|  | ΔSIR_0168::kan^R^ | DUF4176 deletion strain | This study |
|  | ΔSIR_0175::kan^R^ | *essC* deletion strain | (2) |
|  | ΔSIR_1490::kan^R^ | *lapC2* deletion strain | (3) |
|  | ΔSIR_1486-1489::kan^R^ | *telC*, *tipC1*, SIR_1487, *tipC2* deletion strain | (4) |
| *E. coli* XL-1 Blue | *recA1 endA1 gyrA96 thi-1 hsdR17 supE44 relA1 lac* [*F’ proAB lacI^q^* Z Δ M15 Tn*10* (Tet^R^)] | Cloning strain | Agilent |
| *E. coli* BL21 (DE3) CodonPlus | F^-^ *ompT gal dcm lon hsdS*_B_(r_B_^-^ m_B_^-^) λ(DE3) pLysS(Cm^R^) | Protein expression strain | Novagen |

**Table S2: Plasmids used in this study.**

| **Plasmid** | **Relevant features** | **Reference** |
| --- | --- | --- |
| pDL277 | *Streptococcus*-*E. coli* shuttle vector, Spec^R^ | (5) |
| pDL277::P96_*lapA4_*VSV-G | *S. intermedius* expression vector for LapA4, C-terminal VSV-G tag | This study |
| pDL277::P96_*lapA3_*VSV-G | *S. intermedius* expression vector for LapA3, C-terminal VSV-G tag | This study |
| pDL277::P96_SIR_0168*_*VSV-G | *S. intermedius* expression vector for SIR_0168 (DUF4176), C-terminal VSV-G tag | This study |
| pDL277::P96_*lapB4_*VSV-G | *S. intermedius* expression vector for LapB4, C-terminal VSV-G tag | This study |
| pDL277::P96_*lapB3_*VSV-G | *S. intermedius* expression vector for LapB3, C-terminal VSV-G tag | This study |
| pDL277::P96_SIR_0178*_*VSV-G | *S. intermedius* expression vector for SIR_0178 (DUF4176), C-terminal VSV-G tag | This study |
| pDL277::P96_*lapA3_*D79A_VSV-G | *S. intermedius* expression vector for LapA3 D79A mutation, C-terminal VSV-G tag | This study |
| pDL277::P96_*lapA3_*D80A_VSV-G | *S. intermedius* expression vector for LapA3 D80A mutation, C-terminal VSV-G tag | This study |
| pDL277::P96_*telC* | *S. intermedius* expression vector for TelC | (4) |
| pDL277::P96_*telC_*ΔLXG | *S. intermedius* expression vector for TelC LXG domain deletion (TelC_ΔLXG_) | This study |
| pDL277::P96_*sodA*_pep86 | *S. intermedius* expression vector for SodA, C-terminal pep86 tag | This study |
| pDL277::P96_*esxA*_pep86 | *S. intermedius* expression vector for EsxA, C-terminal pep86 tag | This study |
| pDL277::P96_*telA*_LXG_pep86 | *S. intermedius* expression vector for TelA LXG domain (TelA_LXG_), C-terminal pep86 tag | This study |
| pDL277::P96_*telA*_LXG_L41Q_pep86 | *S. intermedius* expression vector for TelA_LXG_ L41Q mutation (TelA_QXG_) C-terminal pep86 tag | This study |
| pDL277::P96_*telC*_LXG_pep86 | *S. intermedius* expression vector for TelC LXG domain (TelC_LXG_), C-terminal pep86 tag | This study |
| pDL277::P96_*telC*_TOX_pep86 | *S. intermedius* expression vector for TelC toxin domain (TelC_TOX_), C-terminal pep86 tag | This study |
| pETDuet-1 | Co-expression vector with *lacI*, T7 promoter, N-terminal His_6_ tag in MCS1, Amp^R^ | Novagen |
| pETduet-1::5′-*lapB4*_flank_SpecPromoter_kanR_3′- *lapB4*_flank::empty | Plasmid containing *S. intermedius* *lapB4* knockout construct for allelic exchange | This study |
| pETduet-1::5′-*lapB3*_flank_SpecPromoter_kanR_3′- *lapB3*_flank::empty | Plasmid containing *S. intermedius* *lapB3* knockout construct for allelic exchange | This study |
| pETduet-1::5′-SIR_0168_flank_SpecPromoter_kanR_3′- SIR_0168_flank::empty | Plasmid containing *S. intermedius* SIR_0168 knockout construct for allelic exchange | This study |
| pETDuet-1::*telA*_LXG__His_6_::*lapA4* | *E. coli* co-expression vector for the TelA LXG domain with LapA4, C-terminal His_6_ on TelA | This study |
| pETDuet-1::*telB*_LXG__His_6_::*lapB4* | *E. coli* co-expression vector for the TelB LXG domain with LapB4, C-terminal His_6_ on TelB | This study |
| pETDuet-1::*telA*_LXG__L41Q_His_6_::*lapA4* | *E. coli* co-expression vector for the TelA LXG domain mutation L41Q with LapA4, C-terminal His_6_ on TelA | This study |
| pET29b | Expression vector with *lacI*, T7 promoter, C-terminal His_6_ tag, Kan^R^ | Novagen |
| pET29b::*lapA3* | *E. coli* expression vector for LapA3 | This study |
| pET29b::*lapB3* | *E. coli* expression vector for LapB3 | This study |
| pET29b::*lapA3_*D79A | *E. coli* expression vector for LapA3 D79A mutation | This study |
| pET29b::*lapA3*_D80A | *E. coli* expression vector for LapA3 D80A mutation | This study |

**Supplemental references**

1.	A. B. Olson *et al.*, Phylogenetic relationship and virulence inference of Streptococcus Anginosus Group: curated annotation and whole-genome comparative analysis support distinct species designation. *BMC genomics* **14**, 895 (2013).

2.	J. C. Whitney *et al.*, A broadly distributed toxin family mediates contact-dependent antagonism between gram-positive bacteria. *eLife* **6** (2017).

3.	T. A. Klein *et al.*, Dual Targeting Factors Are Required for LXG Toxin Export by the Bacterial Type VIIb Secretion System. *mBio* 10.1128/mbio.02137-22, e0213722 (2022).

4.	T. A. Klein, M. Pazos, M. G. Surette, W. Vollmer, J. C. Whitney, Molecular Basis for Immunity Protein Recognition of a Type VII Secretion System Exported Antibacterial Toxin. *Journal of molecular biology* **430**, 4344-4358 (2018).

5.	M. B. Aspiras *et al.*, Expression of green fluorescent protein in Streptococcus gordonii DL1 and its use as a species-specific marker in coadhesion with Streptococcus oralis 34 in saliva-conditioned biofilms in vitro. *Applied and environmental microbiology* **66**, 4074-4083 (2000).
